# PalmLab: A Comprehensive Computational Platform for Systematic Annotation and Functional Interrogation of Protein Palmitoylation

**DOI:** 10.64898/2026.08.30.747671

**Authors:** Shenghong Fu, Wei Wang, Jiaming Huang, Maolin Deng, Qingyuan Li, Zhengqin Rong, Yu-Jian Kang, Bo Xu

## Abstract

Protein palmitoylation is a dynamic and reversible post-translational lipid modification with progressively recognized pathophysiological relevance in cancer and a range of human diseases. Current databases, however, predominantly curate palmitoylation sites derived from isolated literature reports and lack comprehensive cataloging across mass spectrometry (MS)-based palmitoylome datasets, which severely impedes standardized annotation and cross-cohort comparative analyses. Here, we introduce PalmLab, an integrative computational platform that synergistically couples MS-based data curation with interactive, multi-dimensional analytical functionalities for both human and mouse proteomes. Through a unified and rigorously standardized pipeline, PalmLab enables systematic reanalysis of all published palmitoylome datasets, yielding a curated compendium of 15,968 human and 9,924 mouse MS-validated palmitoylated proteins—more than doubling the total number of entries available in existing public repositories. The platform further provides user-friendly, code-free analytical modules that support differential palmitoylation profiling, context-specific pattern visualization, protein-protein correlation network inference, mutation-palmitoylation association mapping, and sequence-based motif discovery. Collectively, PalmLab constitutes a comprehensive and openly accessible resource for systematic data mining and functional interrogation of palmitoylation events, thereby facilitating the elucidation of palmitoylation-mediated regulatory circuits and discovery of novel therapeutic targets. The platform is freely available at https://palmlab.intelligent-oncology.com.

## Introduction

Protein palmitoylation is a lipid post-translational modification catalyzed by the zinc finger DHHC-type containing (ZDHHC) protein family, which critically regulates the localization, stability and function of numerous cellular proteins^1^. Accumulating evidence demonstrates that proteins undergoing palmitoylation are linked to disease pathogenesis and progression^2^, highlighting its potential as a promising therapeutic target for inhibitor development. In cancer, numerous proteins encoded by driver genes were reported to be palmitoylated, which regulate hallmark phenotypes of cancer cells^3^. Given its important role in regulating diverse cellular processes associated with disease progression, particularly cancer, protein palmitoylation is increasingly recognized as an important post-translational modification^4^, highlighting the need for comprehensive resources to facilitate its systematic investigation.

Recent mass spectrometry (MS)-based quantification of palmitoylation enables high-throughput mapping of this modification in different cells and tissues^5^. MS-based palmitoylation proteome (palmitoylome) analysis in hepatocellular carcinoma models systematically uncovered fatty acid synthase (FASN) palmitoylation as a key regulatory mechanism contributing to tumorigenesis^6^. In dendritic cells, large-scale palmitoylome profiling revealed palmitoylation of interferon-induced transmembrane protein 3 (IFITM3) was essential for its antiviral activity^7^. Therefore, MS-based palmitoylome profiling provides a reliable and systematic approach for large-scale identification of palmitoylation sites, quantitative comparison of palmitoylation dynamics across biological contexts and construction of palmitoylation-regulated networks, facilitating the discovery of novel regulatory mechanisms and potential therapeutic targets.

Currently, databases such as SwissPalm^8^, CysModDB^9^, dbPTM^10^ and PTMD^11^ have compiled experimentally validated protein palmitoylation sites from published literature, including those detected by Acyl-Biotin Exchange (ABE) and click-chemistry metabolic labeling with fatty acid analogs, often followed by MS. Although SwissPalm2 incorporates hits from published MS-based palmitoylome studies, these databases primarily curated the proteins or sites reported in the original publications rather than systematically reanalyzing the raw MS datasets using a standardized processing pipeline. Consequently, differences in data processing and reporting across studies limit standardized palmitoylation site annotation and robust comparative analyses. In addition, no open-access platform enables comprehensive cross-study analysis of protein palmitoylation patterns. Thus, we present PalmLab, an integrated resource that provides curated palmitoylated proteins in human and mouse, together with a series of code-free functionalities for palmitoylation pattern analysis (Fig. 1). By uniquely combining standardized MS-based palmitoylation site annotation with novel interactive analysis modules, PalmLab enables systematic exploration of functional palmitoylation events and facilitates the identification of potential regulatory mechanisms and therapeutic targets.

**Fig. 1.**
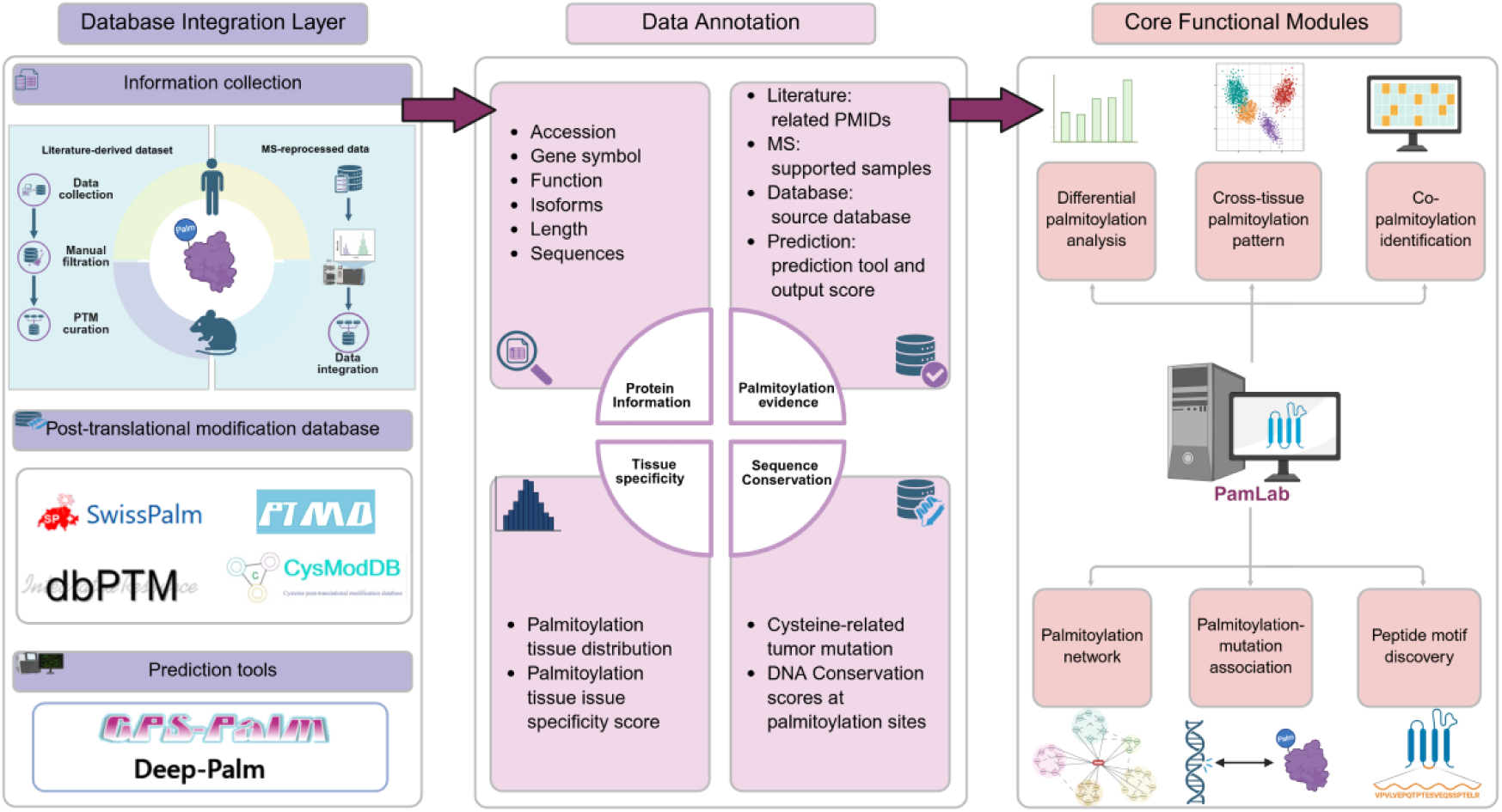
Schema describing the overall architecture of PalmLab.

## Results

### Evidence-Tiered Curation of Protein Palmitoylation Annotations

Accurate annotation of palmitoylated proteins and their modification sites is fundamental to elucidate the underlying molecular mechanisms. Therefore, PalmLab systematically integrates multi-source evidence for palmitoylation site curation in human and mouse. To curate experimentally validated palmitoylated proteins, we manually collected published MS-based palmitoylome studies from PubMed dating back to 2005, representing the broadest coverage among currently available palmitoylation resources (Supplementary Fig. 1 and Supplementary Table 1). By reanalyzing published MS-based palmitoylome datasets using a standardized processing pipeline, PalmLab curated 15,968 human and 9,924 mouse palmitoylated proteins, corresponding to 33,616 and 89,978 modification sites. These MS-supported entries represent approximately twofold greater coverage of the existing databases including SwissPalm, CysModDB, dbPTM and PTMD (Fig. 2A and Supplementary Table 2). In parallel, PalmLab curated the palmitoylated proteins and sites reported in the original publications as literature-based evidence. The standardized MS-derived annotations and literature-based records together comprise the experimentally supported palmitoylation dataset in PalmLab. We further integrated the annotations from the four databases (SwissPalm, CysModDB, dbPTM and PTMD) as database-based evidence. In addition, predicted palmitoylated cysteine sites generated by GPS-Palm^12^ and Deep-Palm^13^ were incorporated as prediction-based evidence. By integrating multi-level evidence from mass spectrometry, literature, databases, and prediction algorithms, PalmLab enables comprehensive and accurate annotation of protein palmitoylation events (Fig. 2B).

**Fig. 2.**
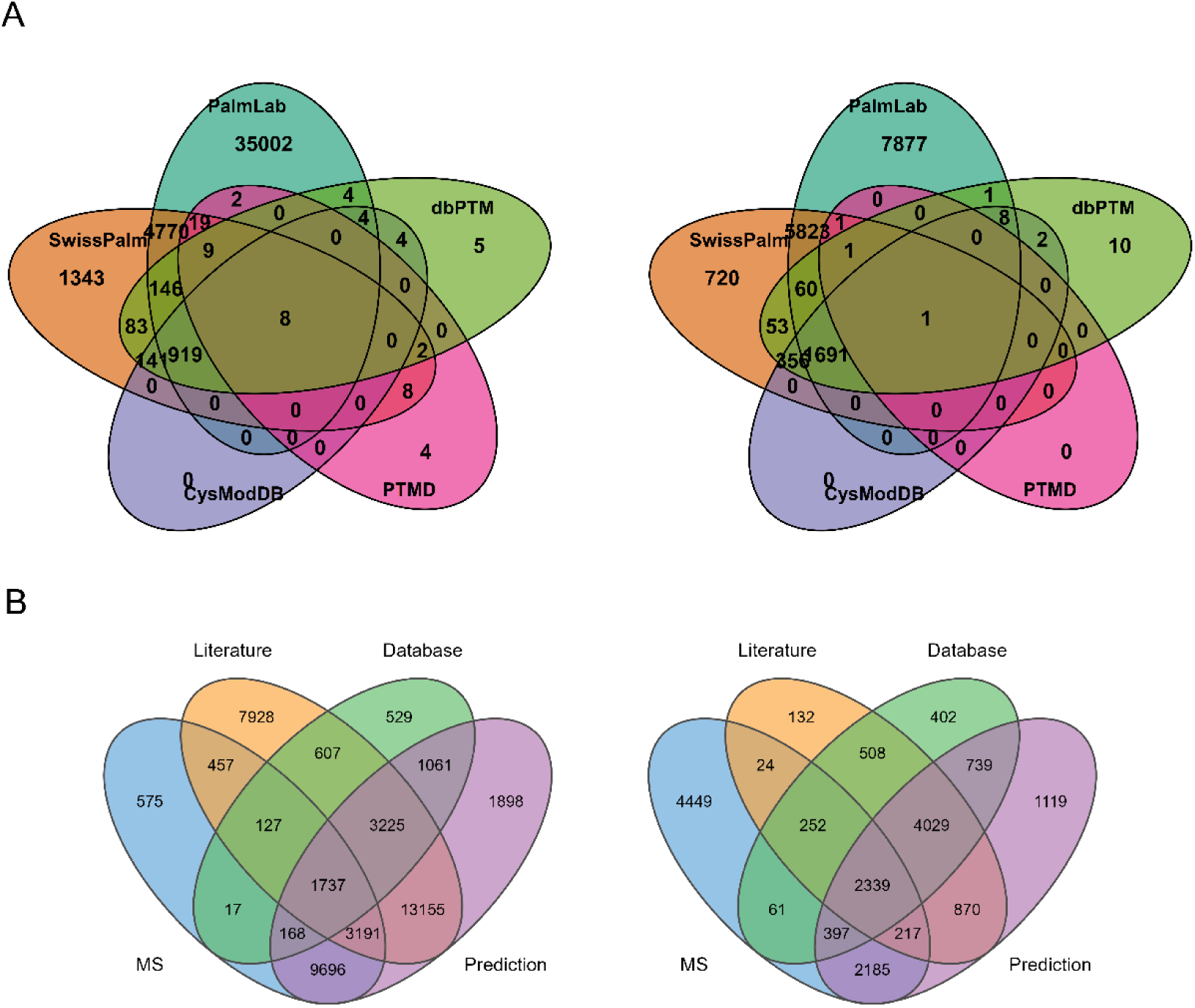
Palmitoylated protein number curated by PalmLab. (A) Comparative coverage of palmitoylated proteins in PalmLab versus existing databases (SwissPalm, CysModDB, dbPTM, and PTMD) in human and mouse. The number of palmitoylated proteins in PalmLab is with experimental evidence (MS-based and literature-based). The left panel shows human data (40883 proteins in PalmLab), and the right panel shows mouse data (15463proteins in PalmLab). (B) Number of palmitoylated proteins from four evidence sources in human and mouse in PalmLab. MS denotes palmitoylated proteins identified through reanalysis of raw mass spectrometry data. Literature represents experimentally reported palmitoylated proteins curated from published studies. Database refers to palmitoylated proteins collected from four public resources, including SwissPalm, CysModDB, dbPTM, and PTMD. Prediction represents palmitoylation predictions generated by GPS-Palm and Deep-Palm. The number of human and mouse proteins are shown in the left and right panels respectively.

### System Architecture and Functional Modules of PalmLab

With the multi-level annotation, PalmLab enables users to query specific genes or proteins to retrieve detailed palmitoylation information (Table 1), including supporting evidence from the primary studies (Fig. 3A), tissue specificity of palmitoylation (Fig. 3B) and mapped modification cysteines (Fig. 3C). Furthermore, since evolutionary conservation at the nucleotide level provides insights into the functional importance of amino acids and their post-translational modifications, PalmLab presents DNA-level conservation scores across vertebrates (PhyloP and PhastCons) for all cysteine residues alongside their palmitoylation status (Fig. 3D). To facilitate the investigation of cancer relevance, we further annotate whether each cysteine residue is affected by non-synonymous somatic mutations identified in tumors from the TCGA cohort (Fig. 3E).

**Table 1.**
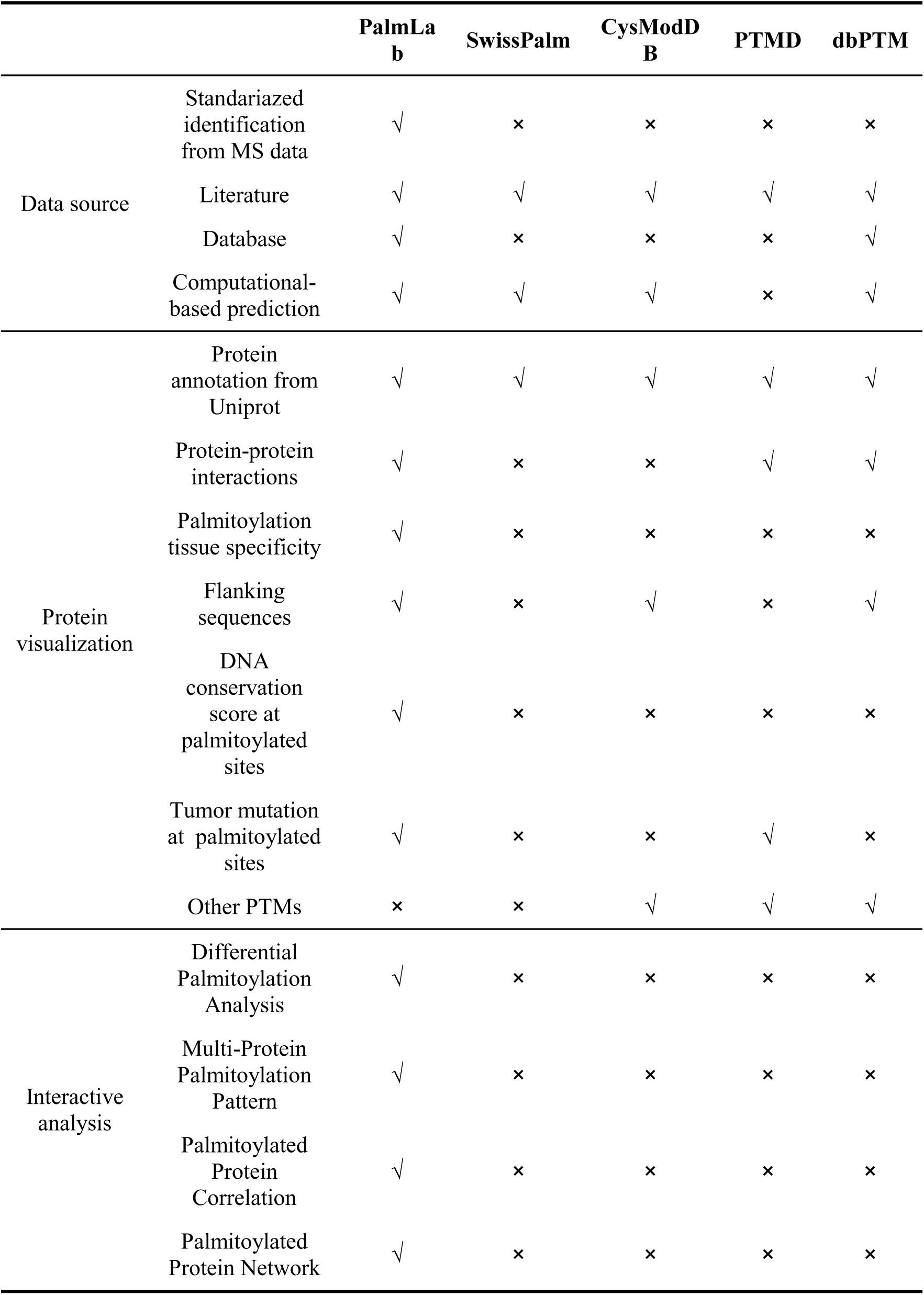
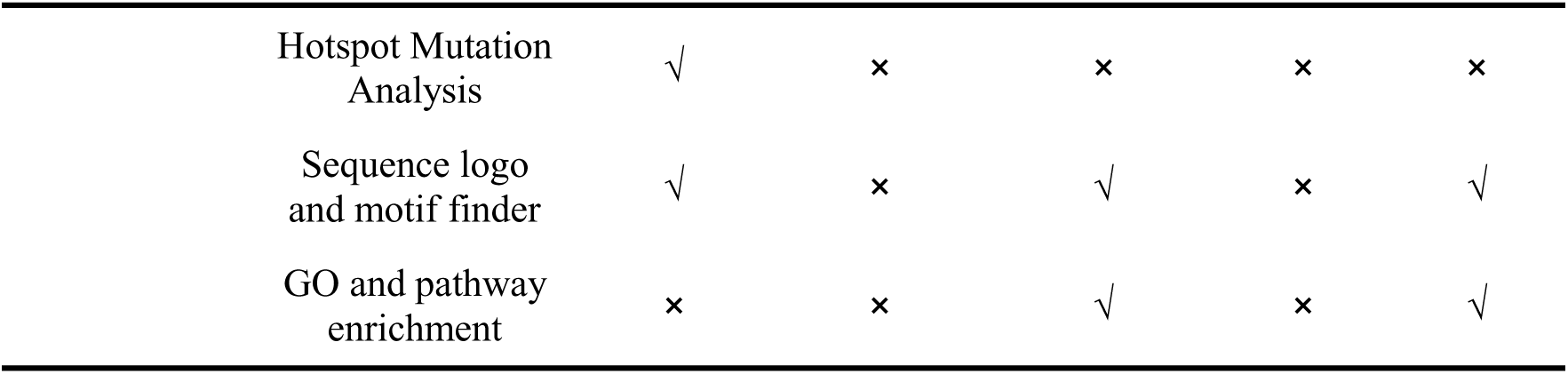
Comparison of PalmLab and existing databases.

|  |  | PalmLab | SwissPalm | CysModDB | PTMD | dbPTM |
| --- | --- | --- | --- | --- | --- | --- |
| Data source | Standardized identification from MS data | √ | × | × | × | × |
|  | Literature | √ | √ | √ | √ | √ |
|  | Database | √ | × | × | × | √ |
|  | Computational-based prediction | √ | √ | √ | × | √ |
| Protein visualization | Protein annotation from Uniprot | √ | √ | √ | √ | √ |
|  | Protein-protein interactions | √ | × | × | √ | √ |
|  | Palmitoylation tissue specificity | √ | × | × | × | × |
|  | Flanking sequences | √ | × | √ | × | √ |
|  | DNA conservation score at palmitoylated sites | √ | × | × | × | × |
|  | Tumor mutation at palmitoylated sites | √ | × | × | √ | × |
|  | Other PTMs | × | × | √ | √ | √ |
| Interactive analysis | Differential Palmitoylation Analysis | √ | × | × | × | × |
|  | Multi-Protein Palmitoylation Pattern | √ | × | × | × | × |
|  | Palmitoylated Protein Correlation | √ | × | × | × | × |
|  | Palmitoylated Protein Network | √ | × | × | × | × |

**Table 1. Comparison of PalmLab and existing databases.**
|  |  |  |  |  |  |
| --- | --- | --- | --- | --- | --- |
| Hotspot Mutation Analysis | √ | × | × | × | × |
| Sequence logo and motif finder | √ | × | √ | × | √ |
| GO and pathway enrichment | × | × | √ | × | √ |

**Fig. 3.**
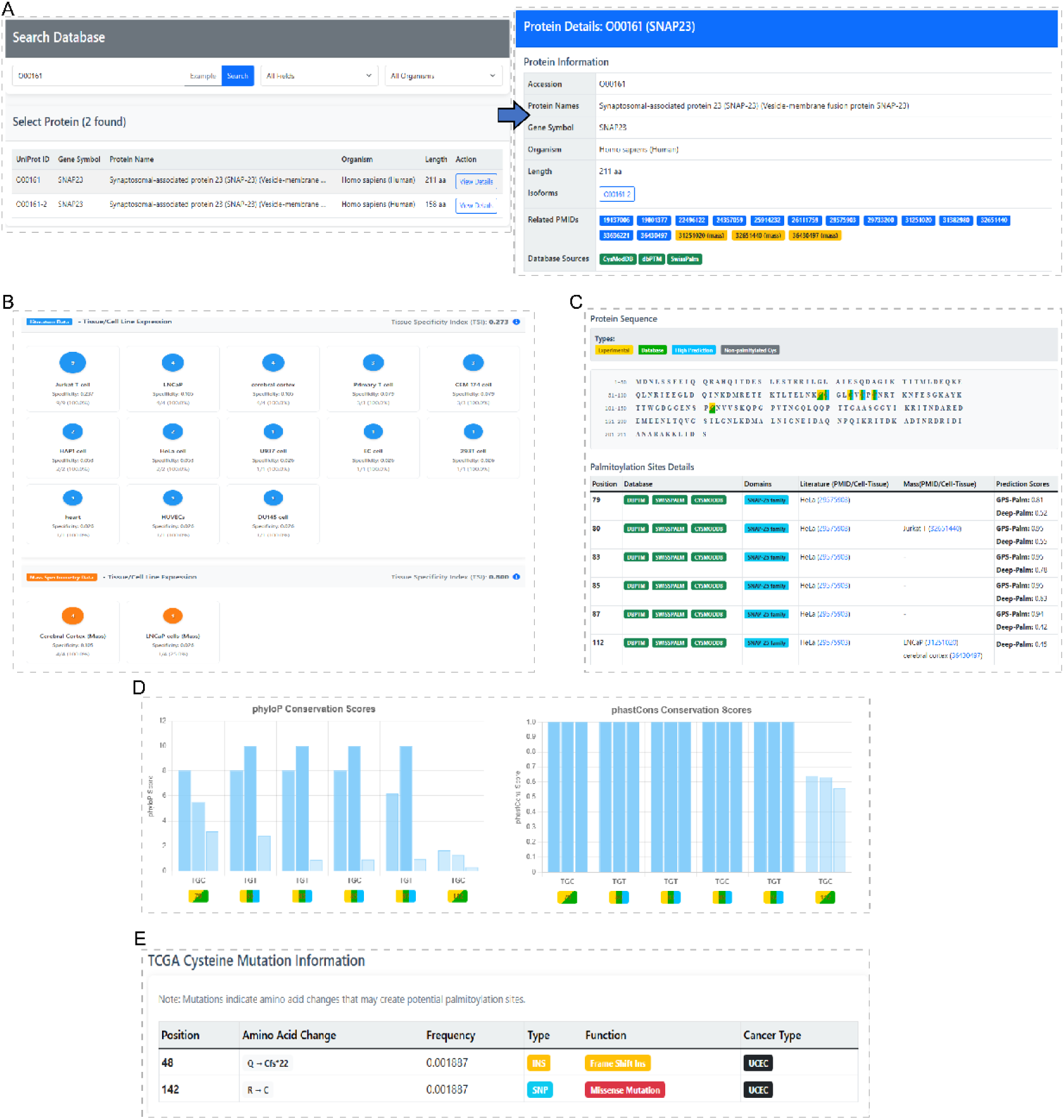
Visualization of protein annotation and palmitoylation information in PalmLab. (A) Basic information of the queried protein. (B) Distribution of protein palmitoylation across different tissues or cell lines. (C) Palmitoylation status of cysteine residues. (D) Conservation scores for the nucleotides encoding cysteine residues. (E) TCGA tumor mutations leading to gain or loss of cysteine residues.

Beyond comprehensive annotation and query functions, PalmLab provides a suite of interactive analysis modules that are not available in existing palmitoylation resources (Table 1). Leveraging standardized palmitoylome datasets, these modules enable users to identify tumor-and tissue-specific palmitoylated proteins, construct protein interaction networks centered on palmitoylated proteins, compare palmitoylation patterns across different tissues, investigate associations between somatic mutations and palmitoylation events, and discover enriched sequence motifs surrounding palmitoylated cysteine residues (Fig. 1). Together, these functionalities allow researchers to move beyond single-site retrieval toward integrative exploration of palmitoylation landscapes and their potential functional relevance.

### Tissue-Specific Profiling of Differential Palmitoylation Patterns

Accumulating evidence has demonstrated that protein palmitoylation exhibits strong context specificity across different tissues, cell types, and pathological conditions^14^. To systematically characterize these context-dependent changes, PalmLab provides functional modules for differential palmitoylation analysis (Fig. 4A) and cross-sample visualization (Fig. 4B), enabling systematic investigation of the dynamic changes in protein palmitoylation across tissues and pathological states. To facilitate biological interpretation, PalmLab pre-integrates curated cancer-related signaling pathways^15^, allowing users to input a list of proteins for systematic palmitoylation profiling and comparative analysis at the pathway level.

**Fig. 4.**
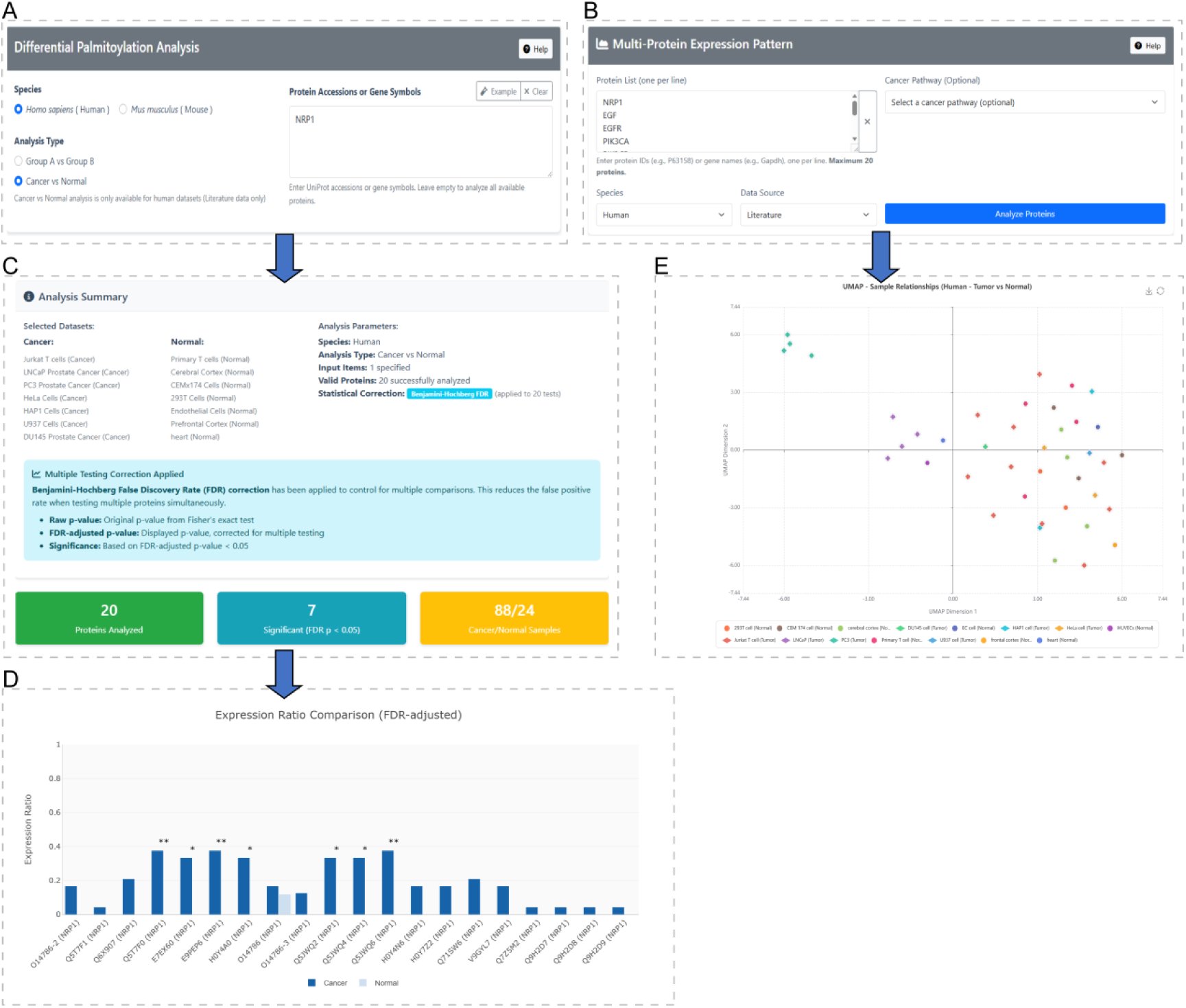
Examples of PalmLab analyses using the “Differential Palmitoylation Analysis” and “Multi-Protein Palmitoylation Pattern” modules. (A) Interface of the “Differential Palmitoylation Analysis” module. (B) Interface of the “Multi-Protein Palmitoylation Pattern” module. (C) Result of searching NRP1 in differential palmitoylation analysis. (D) Barplot to visualize the distribution of palmitoylated sample number in different group.| (E) UMAP visualization based on the palmitoylation profiles of NRP1 and different protein isoforms in the EGF-EGFR-PI3K-AKT signaling network. The input terms include NRP1, EGF, EGFR, PIK3CA, PIK3CB, PIK3CD, AKT1, AKT2, AKT3 and BAD.

To demonstrate the utility of PalmLab for investigating context-specific palmitoylation patterns, we analyzed the palmitoylation profiles of proteins involved in the cancer-related pathways. Using the “Differential Palmitoylation Analysis” module, we identified NRP1, a co-receptor for vascular endothelial growth factor (VEGF) and semaphorin signaling, as a protein exhibiting tumor-specific palmitoylation (Fig. 4C, 4D). NRP1 is a well-established regulator of tumor progression^16–18^ and immune evasion^19,20^ and has been reported to promote prostate cancer progression through activation of the EGFR-dependent PI3K-AKT signaling pathway^21^. To further demonstrate the utility of PalmLab, we used the “Multi-Protein Palmitoylation Pattern” module to visualize the palmitoylation profiles of NRP1 and proteins in the EGF-EGFR-PI3K-AKT signaling network (KEGG entry: N00033) across different tissues and cell types. Among these proteins, NRP1 and EGFR had experimentally supported palmitoylation evidence in PalmLab and exhibited prostate cancer-associated palmitoylation patterns (Supplementary Fig. 2). UMAP dimension reduction based on the palmitoylation profiles of different isoforms in this pathway further revealed a distinct clustering of prostate cancer cell lines (Fig. 4E). To our knowledge, no previous evidence has linked the tumor-promoting function of NRP1 to protein palmitoylation. Using the “Differential Palmitoylation Analysis” and “Multi-Protein Palmitoylation Pattern” modules, PalmLab identified a prostate cancer-associated palmitoylation pattern of NRP1. Together with its well-established oncogenic role^21,22^, this finding highlights NRP1 as a novel candidate for future investigation of palmitoylation-mediated regulation in cancer, while illustrating the utility of PalmLab in uncovering context-specific palmitoylation patterns.

### Construction and Analysis of Palmitoylation-Driven Correlation Networks

Protein palmitoylation functions in a coordinated manner to regulate signaling pathways, membrane trafficking and protein stability^23–26^, resulting in significant co-occurrence (simultaneous palmitoylation of two proteins in the same sample) or mutual exclusion (palmitoylation of the two proteins rarely occurring together in the same sample) relationships. To facilitate the systematic exploration of these relationships, PalmLab provides two complementary analysis modules. The “Palmitoylated Protein Network” module identifies proteins exhibiting significant co-occurrence or mutual exclusion with a query protein and visualizes the resulting associations as an interactive network (Fig. 5A). The “Palmitoylated Protein Correlation” module enables users to directly assess the statistical correlation between any pair of palmitoylated proteins (Fig. 5B).

**Fig. 5.**
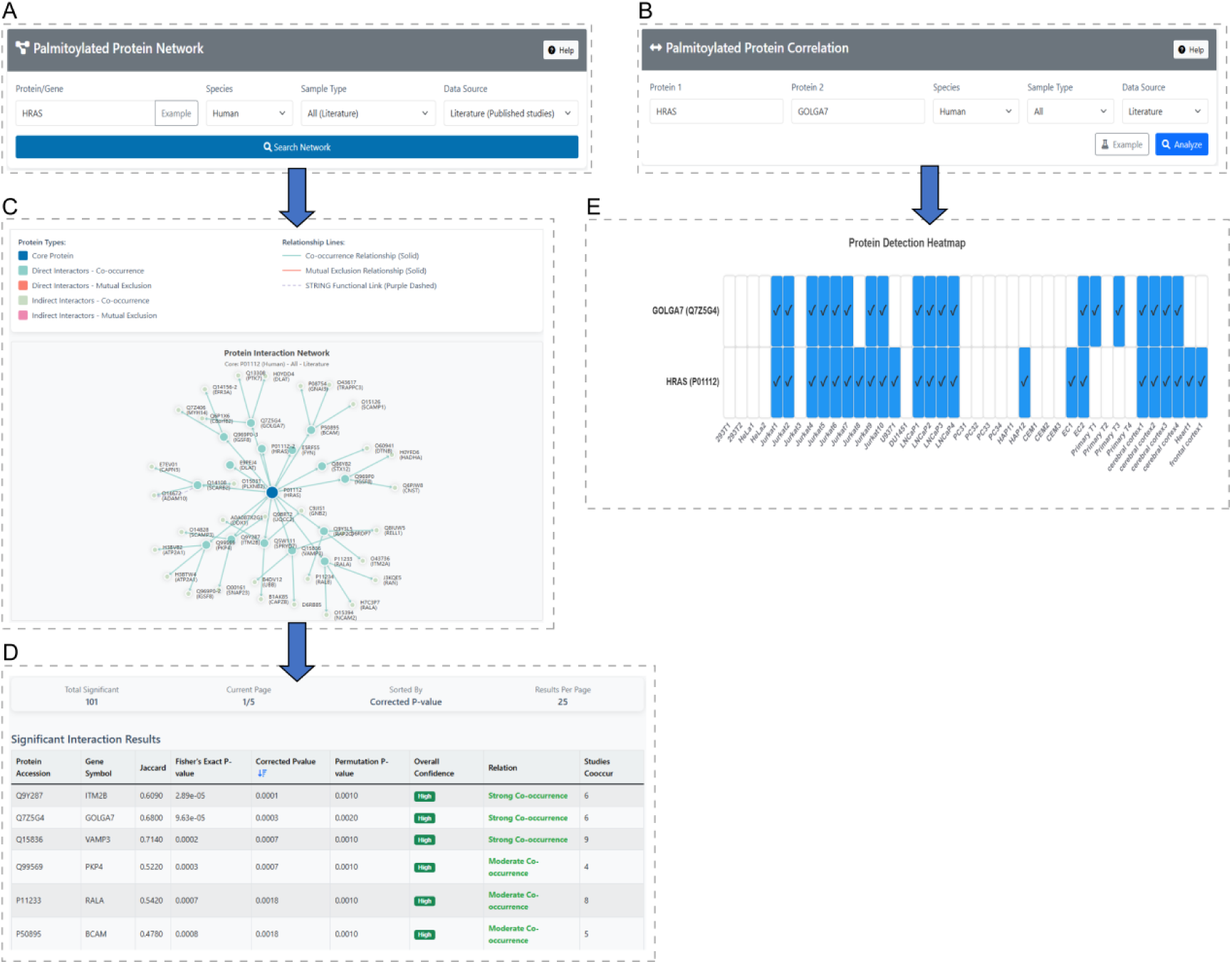
Identification of co-occurring and mutually exclusive palmitoylation relationships using PalmLab. **(A)** Interface of the “Palmitoylated Protein Network” module. **(B)** Interface of the “Palmitoylated Protein Correlation” module. **(C)** HRAS-centered palmitoylation network generated by the “Palmitoylated Protein Network” module, showing proteins with significant co-occurrence or mutual exclusion of palmitoylation with HRAS. **(D)** List of proteins exhibiting the significant palmitoylation associations with HRAS. The most significant associations are with ITM2B and GOLGA7. **(E)** Co-palmitoylation pattern of HRAS and GOLGA7 plotted by “Palmitoylated Protein Correlation” module.

Using the well-characterized oncogenic protein HRAS as an example, the “Palmitoylated Protein Network” module generated an HRAS-centered palmitoylation correlation network, linking proteins that exhibited significant co-occurrence or mutual exclusion of palmitoylation with HRAS (Fig. 5C). Among the identified proteins, ITM2B and GOLGA7 showed the strongest associations with HRAS (Fig. 5D). We then queried HRAS and GOLGA7 in the “Palmitoylated Protein Correlation” module and found that these two proteins exhibited a significant co-palmitoylation pattern across samples (Fig. 5E). GOLGA7 has been reported to stabilize the palmitoyltransferase ZDHHC9, thereby promoting HRAS and NRAS palmitoylation^27,28^. The association identified by PalmLab suggests that GOLGA7 palmitoylation may represent an additional layer of regulation in this process.

### Integrated Analysis of Mutation-Palmitoylation Associations and Sequence Motifs

To facilitate the investigation of the interplay between genomic alterations and protein S-palmitoylation, PalmLab provides a “Hotspot Mutation Analysis” module that systematically evaluates associations between hotspot mutations and protein palmitoylation events across samples (Fig. 6A). The analysis integrates mutation profiles with palmitoylation data in human tumor cells and applies Fisher’s exact test and logistic regression to identify statistically significant mutation-palmitoylation associations (Supplementary Fig. 4). Significant associations are classified as positive or negative correlations and are presented through interactive summary tables (Fig. 6B) and detailed visualization for each pair (Fig. 6C), enabling users to rapidly identify candidate mutation-associated palmitoylation events for further investigation.

**Fig. 6.**
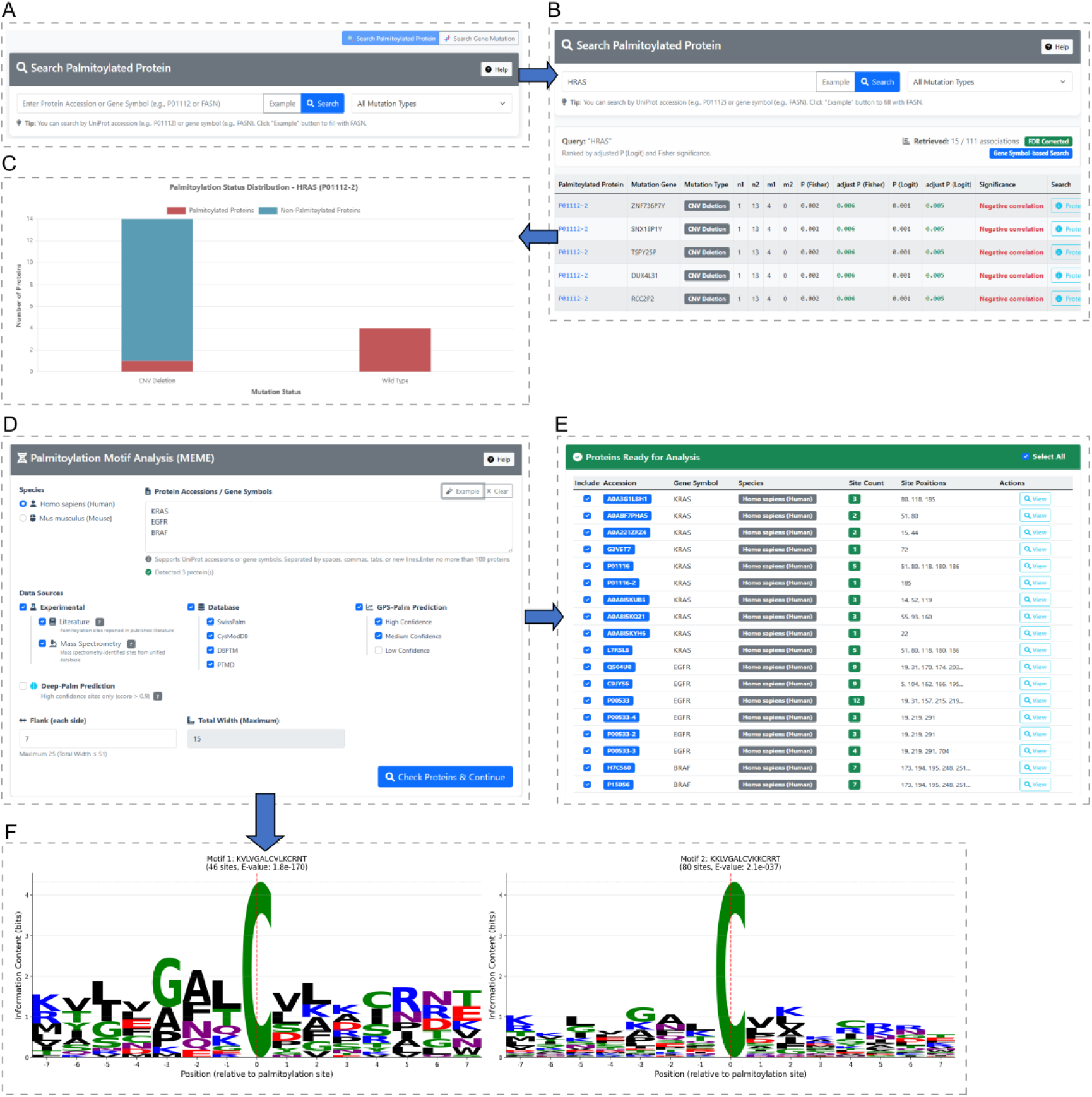
Genomic alteration-associated palmitoylation analysis and motif discovery in PalmLab. **(A)** Interface of the “Hotspot Mutation Analysis” module. **(B)** Summary table displaying significant mutation-palmitoylation associations identified by Fisher’s exact test and logistic regression. **(C)** Detailed visualization of a single hotspot mutation-palmitoylation pair. The two bars compare the numbers of samples with palmitoylated (red) and non-palmitoylated (blue) status in mutant versus wild-type cells. **(D)** Interface of the “Palmitoylation Motif Finder” module. **(E)** Interactive table for selecting palmitoylation sites in query proteins for subsequent motif analysis. **(F)** Identified amino acid sequence motifs surrounding selected palmitoylated cysteines.

PalmLab also incorporates a “Palmitoylation Motif Finder” module for identifying conserved sequence features surrounding palmitoylation sites (Fig. 6D). Users can input a list of proteins and customize the flanking sequence length and evidence level (e.g., experimental, literature-curated, or predicted) for palmitoylation sites. The corresponding amino acid sequences centered on palmitoylated cysteine residues are then extracted for motif enrichment analysis using MEME and visualized as sequence logos with Logomaker (Fig. 6E). Conserved motifs are ranked according to their statistical significance (E-value), allowing users to explore sequence preferences associated with protein palmitoylation and compare motif patterns among different protein sets (Fig. 6F). Together, these novel analysis modules enable genomic-and sequence-level investigations of protein palmitoylation, providing unique capabilities for exploring its regulatory mechanisms.

## Discussions

Protein S-palmitoylation is a reversible lipid post-translational modification that regulates protein localization, trafficking, stability, and signal transduction^4,29,30^. With the increasing application of mass spectrometry-based palmitoylome profiling, large numbers of palmitoylated proteins and modification sites have been identified across diverse biological systems. However, these datasets are distributed across individual studies and generated using heterogeneous experimental protocols and analysis pipelines. To address these limitations, we developed PalmLab, a comprehensive resource that integrates standardized MS-derived palmitoylation data with literature-curated, database-derived and computationally predicted evidence, substantially expanding the coverage of experimentally supported palmitoylated proteins (Fig. 2A). Unlike existing resources that directly curated palmitoylated sites from literature, PalmLab systematically reanalyzes MS raw data using a unified processing pipeline, thereby reducing technical heterogeneity among studies and enabling direct comparison of palmitoylation events across tissues, cell types and diseases.

Beyond serving as a repository of curated palmitoylation events, PalmLab provides a suite of interactive analysis modules designed to facilitate palmitoylome data mining and multi-omics integration for systematic exploration of functional palmitoylation events. Through representative case studies, we demonstrate how these modules can help researchers uncover context-specific palmitoylation patterns and identify unrecognized regulatory relationships. For example, PalmLab identified a prostate cancer-associated palmitoylation pattern of NRP1, a well-established regulator of tumor progression and immune evasion for which no previous evidence of palmitoylation has been reported. In addition, PalmLab revealed a significant association between GOLGA7 and HRAS palmitoylation. Given that GOLGA7 is an essential cofactor of the palmitoyltransferase ZDHHC9 required for HRAS palmitoylation, this finding generates a new hypothesis that GOLGA7 palmitoylation itself may participate in this pathway.

In summary, PalmLab provides a comprehensive resource that combines standardized MS-based protein palmitoylation curation with multi-level evidence integration and a suite of novel analysis tools for systematic palmitoylome exploration. By combining large-scale data collection, standardized annotation and user-friendly downstream analyses, PalmLab offers a valuable platform for systematic investigation of protein palmitoylation and facilitates the discovery of context-dependent regulatory mechanisms and potential therapeutic targets.

## Materials and Methods

### Web Server Architecture

All data are processed and organized within a PostgreSQL database management system. The backend of the PalmLab platform is built on PostgreSQL 14.22 (database server). The web server is deployed on an Ubuntu 22.04.5 LTS virtual machine, using Nginx 1.18.0 as the HTTP server and Gunicorn 25.3.0 as the WSGI application server. The web interface is developed using the Django template engine 5.1. For frontend data visualization, PalmLab integrates ECharts (version 5.4.3 on analysis pages and version 5.5.0 on the home page) and Chart.js 4.4.0 for interactive statistical charting and visualization. The user interface is designed using Bootstrap 5.1.3 to ensure responsive layout compatibility across desktop and mobile devices. The PalmLab platform has been tested across major web browsers, including Google Chrome, Mozilla Firefox, Apple Safari and Microsoft Edge, to ensure cross-platform compatibility and consistent user experience. The PalmLab web server is accessible at https://palmlab.intelligent-oncology.com/.

### Data Collection

To construct a reliable palmitoylome dataset, we systematically retrieved published studies of palmitoylated protein identification via mass spectrometry in PubMed since 2005 with the following queries: **a)** (“Palmitoylation” OR “Acylation”) AND (“Mass Spectrometry” OR “LC–MS/MS”) AND (“Enrichment” OR “Proteomics”) AND (“Human” OR “Mouse”) or **b)** “protein palmitoylation” AND “mass spectrometry”. After manual screening for relevance and eligibility, the identified datasets were subjected to a rigorous standardization procedure (Supplementary Table 1). The palmitoylated proteins and sites reported by these studies were integrated into the PalmLab resource and curated as literature-supported entries. Other data resources incorporated in PalmLab are provided in Supplementary Table 3.

### MS Data Processing

For studies with raw mass spectrometry data, we searched against UniProt Human or Mouse reference proteomes using MaxQuant 2.7.5. For each study, we manually curated the original experimental information, including enrichment strategy, mass spectrometer, modifications and search parameters. Based on the most commonly adopted settings across the collected studies and considering the compatibility of different MS workflows, we established a standardized search strategy for uniform reanalysis while incorporating study-specific modifications when required by the original experimental design. Palmitoyl (C), Carbamidomethyl (C), Oxidation (M), Acetyl (Protein N-term) and Deamidation (NQ) were set as variable modifications for all samples. For two datasets (PMID: 34884899 and 29217618), Methylthio (C) was additionally included as a variable modification. The modification for identified palmitoylation sites were described in Supplementary Table 4. The precursor mass tolerance was set to 20 ppm for the first search and 4.5 ppm for the main search, and fragment mass tolerances were set to 20 ppm and 0.5 Da for FT and IT detectors, respectively. A maximum of two missed cleavages was allowed and the minimum peptide length was set to 7 amino acids. For the identification of palmitoylated peptides, a 1% false discovery rate (FDR) was applied at the PSM, protein and site levels with a minimum Andromeda score of 40 and a minimum delta score of 6. This reprocessing generated a standardized, high-confidence dataset of MS-based palmitoylated proteins and sites. All proteins were annotated based on the UniProtKB protein accession IDs.

### Differential Palmitoylation Analysis

PalmLab provides a differential palmitoylation analysis module to enable comparison of palmitoylation modification levels of target proteins across different samples. The statistical significance of differences in palmitoylation status between two sample groups was calculated by Fisher’s exact test. Multiple testing corrections were performed using the Benjamini-Hochberg (BH) method to control the FDR. The statistical significance of the results is indicated by the BH-adjusted p-value and marked with asterisks denoting significance levels (*FDR < 0.05, **FDR < 0.01, ***FDR < 0.001).

### Palmitoylated Protein Correlation Calculation

To systematically identify associations among palmitoylated proteins, stringent data quality control criteria were first applied (Supplementary Fig. 3). Only proteins detected in at least 20% of total samples and present in at least two independent studies were retained. For reproducible protein pair, we required that its co–occurrence or mutual–exclusion pattern could be observed across at least two studies.

For each protein pair, a 2×2 contingency table was constructed from the numbers of samples in which both proteins, either protein alone, or neither protein were detected, as previously described by Li *et al*^31^. Fisher’s exact test was used to calculate the odds ratio (OR) and corresponding p-value, followed by BH correction. The Jaccard index was additionally calculated to quantify the degree of co-occurrence between protein pairs. We classified protein–protein pairs with the following criteria: a) co–occurrence: FDR < 0.05 and OR > 1, b) mutual exclusion: FDR < 0.05 and OR < 1, c) independent: FDR ≥ 0.05 or OR = 1. The Jaccard index was used to stratify co-occurrence pairs’ association strength levels: strong (≥0.6), moderate (0.3-0.6) and weak (<0.3).

To further reduce false-positive associations arising from sample distribution, permutation tests were performed for all significant protein pairs (FDR < 0.05). For each permutation, the occurrence frequency of each protein was preserved while its sample labels were randomly shuffled, thereby maintaining the marginal distribution of each protein. Fisher’s exact test was recalculated after each of 1,000 permutations to generate a null distribution of association statistics. Empirical p-values were calculated as the proportion of permuted statistics equal to or more extreme than the observed statistic, and only protein pairs with an empirical p-value < 0.05 were labeled as high-confidence associations in the database.

### Cancer Genomic Alteration-Associated Palmitoylation Identification

To enable systematic investigation of the associations between protein palmitoylation and tumor genomic alterations (CNVs, SNVs and hotspot mutated genes), PalmLab has developed a module for hotspot mutation-associated palmitoylation identification (Supplementary Fig. 4). For mutation-palmitoylation association analysis, we first matched cell lines profiled in the MS-based palmitoylome datasets with corresponding cell lines from the Cancer Cell Line Encyclopedia (CCLE). For each gene, CNV/SNV and gene alteration status from CCLE (Supplementary Table 3) were integrated with the palmitoylation status of the corresponding protein in matched cell lines. Mutations detected in only one cell line were excluded and only proteins detected as palmitoylated in at least 10% of the analyzed samples and supported by at least two independent studies were considered to reduce potential biases from rare events.

For each protein-mutation pair, a 2×2 contingency table was constructed based on the presence or absence of genomic alterations and protein palmitoylation across samples. Fisher’s exact test was performed to evaluate the statistical association between mutation status and palmitoylation status. Firth logistic regression^32,33^ analysis was further performed with palmitoylation status as the binary outcome variable, mutation status as the primary explanatory variable, and cell line identity as a covariate to control for cell line-specific genetic backgrounds. Associations were considered significant when the regression coefficient of mutation status was statistically significant. Multiple testing correction was performed using the BH correction for both Fisher test and logistic regression. Associations were classified as “Positive correlation”, “Negative correlation” or “Not significant” according to the sign of the logistic regression coefficient and the adjusted p-values (significant pairs were defined as adjusted p-values < 0.05 in both Fisher test and logistic regression).

### Amino Acid Sequence Motif Scanning Around Palmitoylated Sites

The “Palmitoylation Motif Finder” module first verifies the input protein identifiers or gene symbols. It then retrieves all annotated palmitoylation sites for the selected proteins, including sites supported by experimental evidence, curated databases and computational predictions. Users can view the details of each palmitoylation site and select the subset of peptides for subsequent motif analysis. PalmLab extracts peptide sequences centred on palmitoylation cysteines with fixed flanking length specified by the user for *de novo* motif discovery using MEME^34^. MEME identifies significant over-represented sequence motifs using the expectation-maximization (EM) algorithm and visualizes them as sequence logos, in which the height of each amino acid reflects its positional frequency. For each identified motif, PalmLab records the corresponding palmitoylation sites and presents the associated peptide sequences in a searchable table, with amino acids color-coded according to their physicochemical properties, allowing users to trace the exact sequences contributing to each motif.

## Contributions

B.X. and Y.K. designed the project. J.H. designed the user interface and built the server base system; S.F., W.W. and Q.L. collected the data and developed the analysis functionalities; M.D. calculated the palmitoylation site prediction results; Y.K., B.X. and S.F. wrote the manuscript. Z.R. advised on the overall architecture, security and maintenance for the website.

## Data availability

PalmLab is publicly accessible at https://palmlab.intelligent-oncology.com. The data and analysis results could be accessed through the API. This website is free and open to all users and there is no login requirement.

