## Supplementary Materials for "PalmLab: A Comprehensive Computational Platform for Systematic Annotation and Functional Interrogation of Protein Palmitoylation"

| PMID | Sample source | Year | Number of samples | High-confidence cutoff |
|---|---|---|---|---|
| Homo sapiens, Literature-derived | | | | |
| 26111759 | Jurkat T cells* | 2015 | 6 | H/L ratio ≥ 1.75 |
| | T cells | | 4 | H/L ratio ≥ 3.0 |
| 31382980 | U937* | 2019 | 1 | - |
| 25914232 | CEMx174 | 2015 | 3 | - |
| 22496122 | Endothelial cells | 2012 | 1 | $p < 0.05$; Mascot score (+HA) > 56; Mascot score (−HA) < 56 |
| 29733200 | HAP1* | 2018 | 2 | PSMs ≥ 3; Average ratio > 2 |
| | HEK293T | | 2 | |
| 19801377 | DU145* | 2010 | 1 | Protein level: $p < 0.05$ (EXP/CON > 6.7);<br>Peptide level: $p < 0.05$ (EXP/CON > 2.4) |
| 37611173 | HeLa* | 2023 | 1 | - |
| 26865113 | SW480 | 2016 | 1 | +$NH_2OH$/−$NH_2OH$ > 2 |
| 26876311 | Frontal cortex | 2016 | 1 | - |
| 24357059 | HUVECs | 2014 | 1 | SILAC ratio (H/L) > 1.5; Peptides ≥ 2 |
| 21076176 | Jurkat T cells* | 2011 | 1 | Normalized peptide counts > 1.5-fold (vs control) |
| 19137006 | Jurkat T cells* | 2009 | 2 | (i) Average spectral count ≥ 5;<br>(ii) Detected in ≥ 3 replicates;<br>(iii) Spectral count ratio ≥ 5 (17-ODYA vs control) |
| 29575903 | HeLa* | 2018 | 2 | Chemical reporter: ≥ 5-fold MS/MS counts (alk-16 vs DMSO or alk-16+$NH_2OH$ vs DMSO);<br>Acyl-RAC: ≥ 5-fold MS/MS counts (+$NH_2OH$ vs −$NH_2OH$) |
| 31251020 | LNCaP* | 2019 | 4 | $q < 0.01$; log2($Hyd^+/Hyd^-$) > 1; ratio ≥ 1.5 |
| 32944167 | PC3* | 2020 | 4 | FDR < 0.05; normalized expression difference ≥ 0.1 |
| 32651440 | Jurkat T cells* | 2020 | 1 | $p < 0.05$ |
| 36430497 | Cerebral cortex | 2022 | 4 | - |
| 33636221 | Heart | 2021 | 1 | PSM ≥ 2 (+hydroxylamine(HxA));<br>Detected in +HxA only or PSM ratio ≥ 2 (+HxA/−HxA) |
| Mus musculus, Literature-derived | | | | |
| 34884899 | Liver | 2021 | 2 | $p < 0.05$; ratio ≥ 1.5 |
| 34200797 | Brain (C57BL/6J) | 2021 | 2 | $p < 0.05$; FDR < 0.01 |
| 35358180 | Brain | 2022 | 1 | $p < 0.05$ |
| 31772009 | NSCs | 2019 | 1 | log2(fold change) > 2 |

| | | | | |
|---|---|---|---|---|
| 31311849 | Brain | 2019 | 1 | p < 0.05; FDR < 0.01 |
| 28526873 | Liver | 2017 | 1 | score > 25 |
| 37925639 | Liver | 2023 | 3 | p < 0.05 |
| 35477839 | Testis | 2022 | 2 | - |
| 29660268 | Brain | 2018 | 1 | protein FDR ≤ 2%; unique peptides ≥ 2 |
| 29217618 | RAW264.7 | 2018 | 1 | p ≤ 0.05; ln(ratio) ≥ 2.5 |
| 39008777 | Lung epithelial cells (MLE12) | 2024 | 1 | log2(fold change) > 0.59; FDR < 0.05 |
| | Brain (C57BL/6) | | 1 | |
| 28680068 | Forebrain | 2017 | 1 | FDR ≤ 0.05; (HA+/HA−) > 3 |
| 26165157 | Liver (C57BL/6J) | 2015 | 1 | - |
| 29733200 | Brain | 2018 | 1 | PSMs ≥ 3 |
| 29575903 | RAW264.7 | 2018 | 3 | Chemical reporter: ≥ 5-fold MS/MS counts (alk-16 vs DMSO or alk-16+$NH_2OH$ vs DMSO);<br>Acyl-RAC: ≥ 5-fold MS/MS counts (+$NH_2OH$ vs −$NH_2OH$) |
| Homo sapiens, MS-based proteomics | | | | |
| 31251020 | LNCaP | 2019 | 4 | log2(Hyd+/Hyd−) > 1 |
| 32651440 | Jurkat T cells | 2020 | 10 | log2(Hyd+/Hyd−) > 1 |
| 36430497 | Cerebral cortex | 2022 | 4 | - |
| Mus musculus, MS-based proteomics | | | | |
| 34884899 | Liver | 2021 | 2 | log2(Hyd+/Hyd−) > 1 |
| 31772009 | NSCs | 2019 | 1 | log2(Hyd+/Hyd−) > 1 |
| 31311849 | Brain | 2019 | 6 | - |
| 37925639 | Liver | 2023 | 6 | - |
| 29217618 | RAW264.7 | 2018 | 10 | - |
| 35358180 | Brain | 2022 | 50 | - |

Supplementary Table 1. Sample annotation.
"*" indicates tumor cell lines from the CCLE dataset used in the "Hotspot Mutation Analysis" module.

**Comparison of datasets incorporated in existing databases**

To date, SwissPalm is the only database that has utilized mass spectrometry-derived data without reanalysis. SwissPalm collected a total of 50 MS-based palmitoylation studies, including 21 studies conducted under non-wild-type experimental conditions (gene knockout, overexpression, or drug treatment); these were subsequently excluded by PalmLab. The remaining 29 studies were incorporated, along with two additional studies uniquely curated by PalmLab (PMID: 31382980, PMID: 39008777).

| | | PalmLab (MS only) | SwissPalm | CysModDB | PTMD | dbPTM |
|---|---|---|---|---|---|---|
| Number of palmitoylated proteins with experimental evidence | Human | 15968 | 7448 | 1076 | 52 | 1326 |
| | Mouse | 9924 | 8706 | 2058 | 3 | 2183 |
| Number of palmitoylation sites with experimental evidence | Human | 28443 | 2874 | 1660 | 32 | 2144 |
| | Mouse | 30531 | 4287 | 3973 | 0 | 3929 |

Supplementary Table 2. Number of palmitoylated proteins with MS evidencen in PalmLab and total number of proteins in exsisting databases.

| Data set | Resource | Website |
|---|---|---|
| Protein palmitoylation experimental evidence | MS-based palmitoylome studies retrieved from PubMed | Supplementary Table 1 |
| Protein palmitoylation sites annotation in databases | SwissPalm | https://swisspalm.org/ |
| | dbPTM | https://biomics.lab.nycu.edu.tw/dbPTM/ |
| | PTMD | https://ptmd.biocuckoo.cn/ |
| | CysModDB | https://cysmoddb.bioinfogo.org/ |
| Genome conservation score | phyloP vertebrates 35way<br>phyloP vertebrates 100way<br>phastCons vertebrates 35way<br>phastCons vertebrates 100way | https://genome-euro.ucsc.edu/cgi-bin/hgTables |
| Patient somatic mutations | TCGA | https://portal.gdc.cancer.gov/ |
| Cancer cell mutations | CCLE OmicsCNGene | https://depmap.org/portal/data_page/?tab=allData&releasename=DepMap%20Public%2024 |

| | | |
|---|---|---|
| | | Q4&filename=OmicsCNGene.csv |
| | CCLE OmicsSomaticMutations | https://depmap.org/portal/data_page/?tab=allData&releasename=DepMap%20Public%2024Q4&filename=OmicsSomaticMutations.csv |
| | CCLE OmicsSomaticMutationsMatrixHotspot | https://depmap.org/portal/data_page/?tab=allData&releasename=DepMap%20Public%2025Q2&filename=OmicsSomaticMutationsMatrixHotspot.csv |
| Protein-protein interactions | STRING | https://cn.string-db.org/cgi/ |

Supplementary Table 3. Public data sets used in PalmLab.

| PMID | Sample source | Modifications |
| --- | --- | --- |
| Homo sapiens, MS-based proteomics | | |
| 31251020 | LNCaP | Nethylmaleimide (C) |
| 32651440 | Jurkat T cells | Nethylmaleimide (C) |
| 36430497 | Cerebral cortex | Carbamidomethyl (C) |
| Mus musculus, MS-based proteomics | | |
| 34884899 | Liver | Methylthio (C) |
| 31772009 | NSCs | Nethylmaleimide (C) |
| 31311849 | Brain | Carbamidomethyl (C) |
| 37925639 | Liver | Carbamidomethyl (C) |
| 29217618 | RAW264.7 | Methylthio (C) |
| 35358180 | Brain | Carbamidomethyl (C) |

Supplementary Table 4. Modifications in Site Analysis.

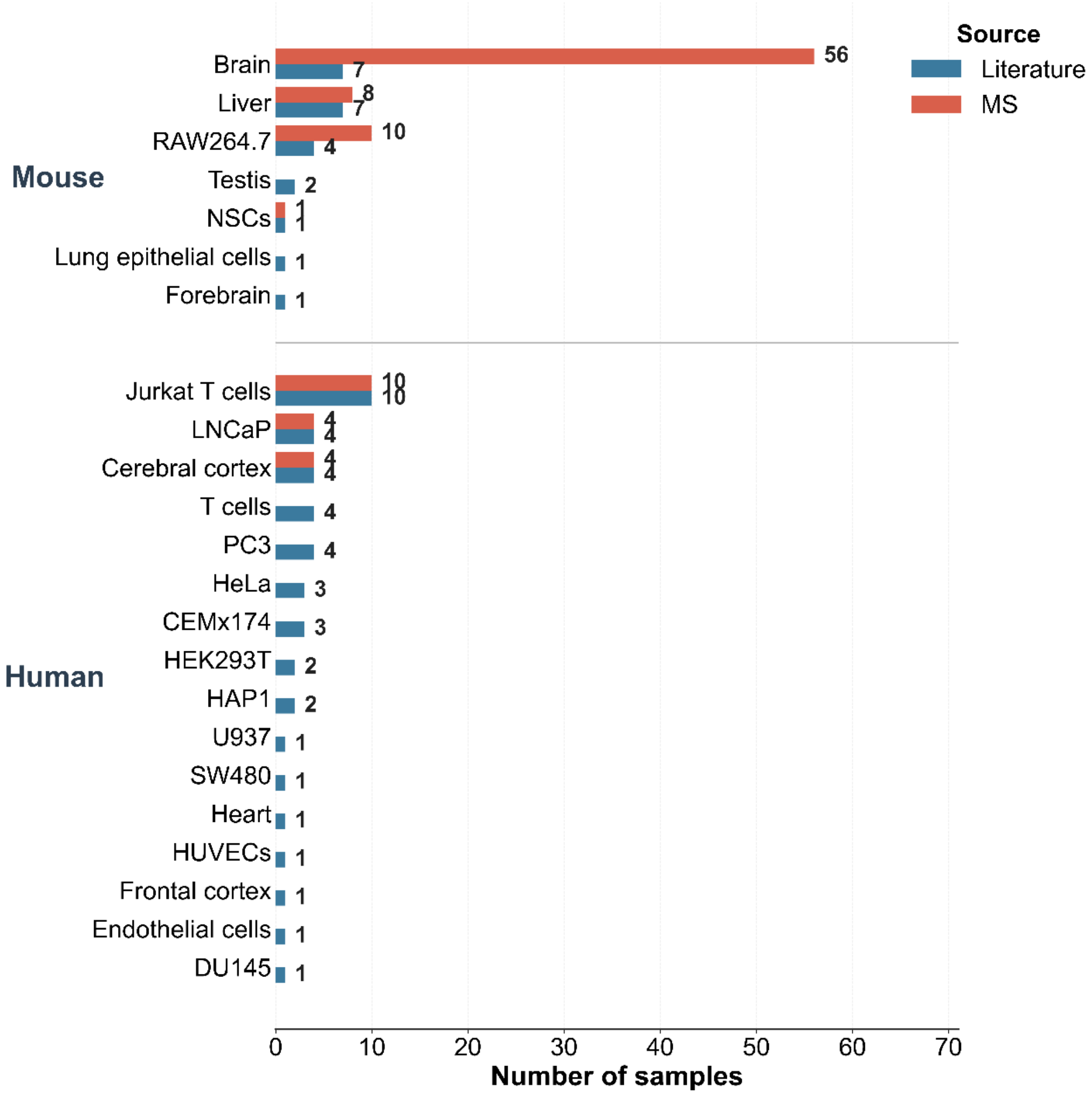


Supplementary Fig. 1. Distribution of sample species and data sources.

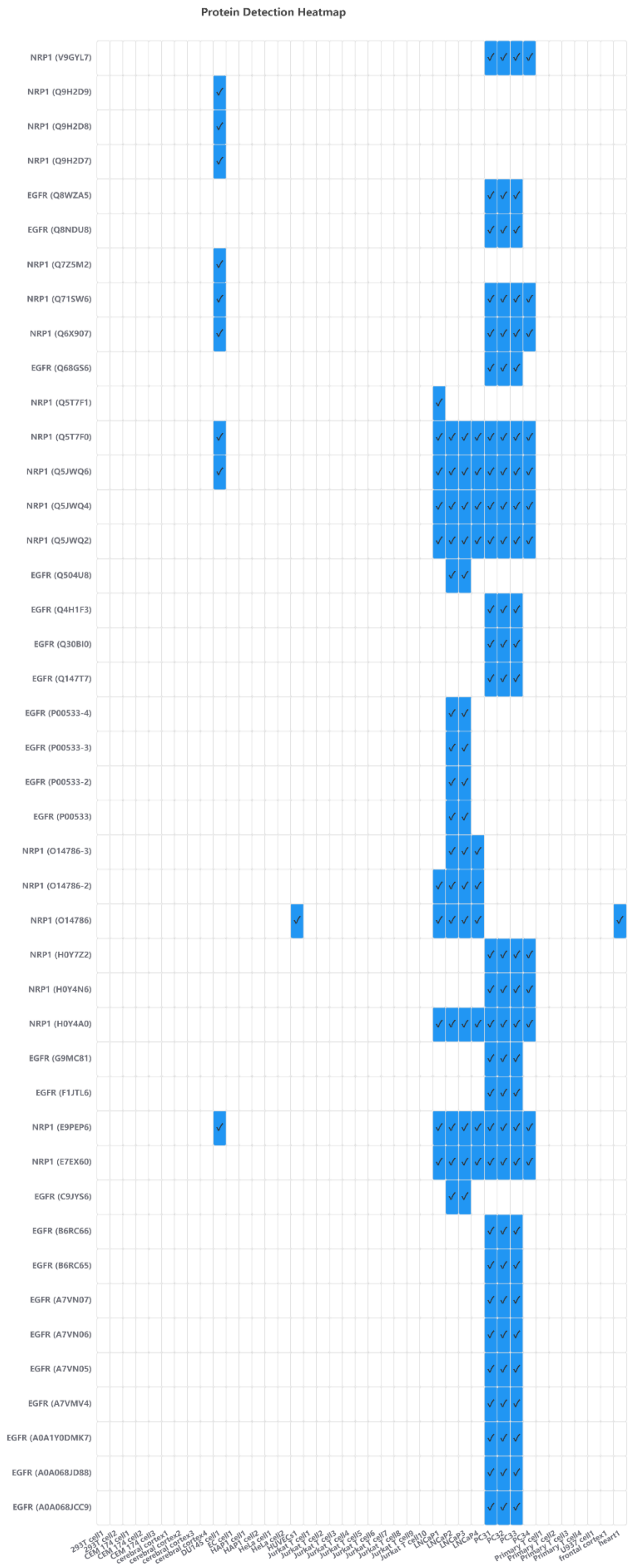


Supplementary Fig. 2. Heatmap visualization of the palmitoylation profiles of NRP1 and protein isoforms in the EGF–EGFR–PI3K–AKT signaling network.

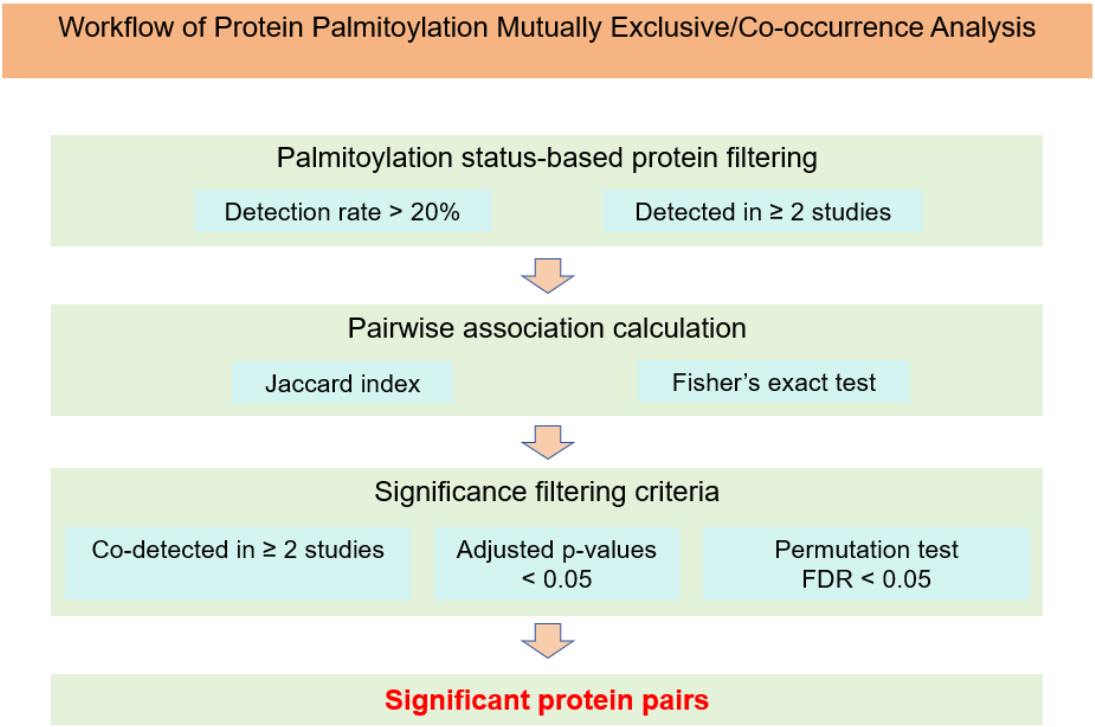


Supplementary Fig. 3. Workflow of Protein Palmitoylation Mutually Exclusive/Co-occurrence Analysis.

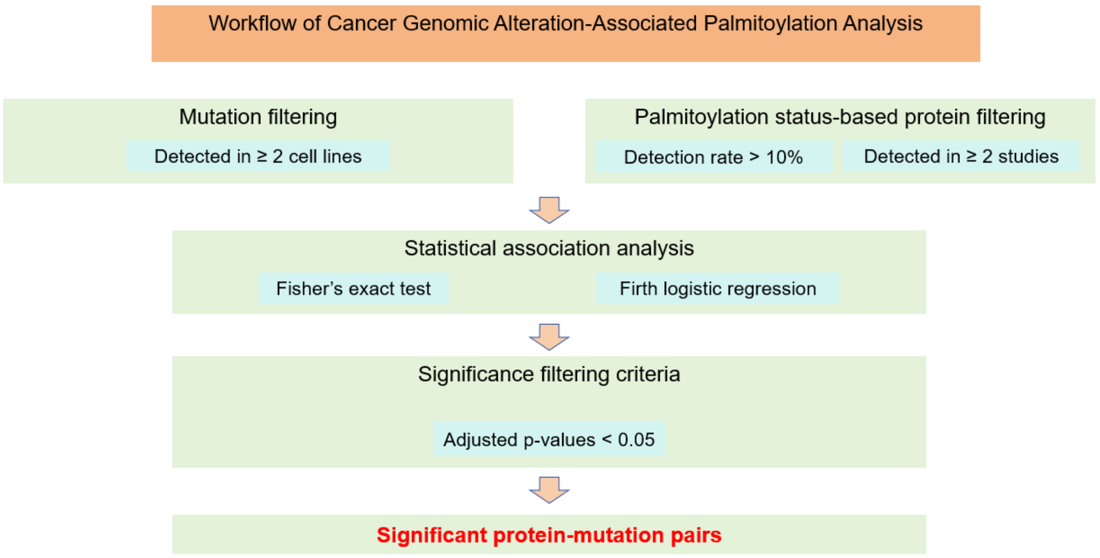


Supplementary Fig. 4. Workflow of Cancer Genomic Alteration-Associated Palmitoylation Analysis.
